# Cerebrovascular Structure–Flow Gradients Revealed by Volumetric Ultrasound Localization Microscopy

**DOI:** 10.64898/2026.10.27.762274

**Authors:** Jingwen Zhu, Shusei Kawara, Kai Riemer, Bingxue Wang, Jipeng Yan, Sophie V. Morse, Simon R. Schultz, James J. Choi, Meng-Xing Tang

**Affiliations:** Department of Bioengineering, Imperial College London, London SW7 2AZ, UK; Charité Center for Biomedicine, Charité - Universitätsmedizin Berlin, Berlin 10117, Germany; Harbin Institute of Technology, Heilongjiang 150001, China; UK Dementia Research Institute Centre, Imperial College London, London W12 0BZ, UK

**Keywords:** mouse brain, volumetric ultrasound localization microscopy, cerebrovasculature, gradients, brain organization

## Abstract

Characterizing the joint organization of microscale cerebrovascular structure and flow in vivo is challenging. Volumetric ultrasound localization microscopy (ULM) provides spatially resolved measures of microvascular structure and flow from the same acquisition, but how these features are jointly organized across brain networks remains unclear. Here, we characterized local vascular phenotypes from transcranial volumetric ULM in mice and examined whether they contained reproducible low-dimensional organization. Local measures of vascular structure and flow dynamics were summarized within corresponding brain regions across mice. Subject-specific vascular phenotype similarity matrices were combined into a group-consensus graph and analyzed using diffusion embedding. Two leading gradients, showing distinct spatial distributions across brain networks, were reproducibly recovered across subject and spatial scales. The principal gradient was associated with local vascular density and flow complexity, defining a vascular structure-related axis. The secondary gradient was associated with flow speed and acceleration, defining a flow-dynamics-related axis. These gradients provide a low-dimensional representation of the cerebrovascular structure and flow and provide a reference for studying coordinated vascular changes across disease, aging, and intervention.

## 01 Introduction

The brain is the most energy-demanding organ in the body. It relies on the cerebrovascular system for continuous delivery of metabolic substrates and for removal of metabolic waste [1,2]. Consequently, vascular organization provides a structural foundation for brain function, and cerebrovascular abnormalities are closely linked to various brain pathologies [3]. Mouse models provide an accessible system for investigating this organization and its alteration in disease [4,5]. Early whole-brain studies using MRI, micro CT [6], and optical microscopy [7,8] have mapped major vessels and revealed marked regional variation in microvascular architecture [6–10]. As they are largely ex vivo studies, the joint in vivo organization of local vascular structure and flow across the mouse brain remains underexplored.

Recently, volumetric ULM has shown the ability to noninvasively characterize whole-brain vascular structure and flow dynamics in small animals, providing a faithful representation of in vivo vascular organization [11–13]. These ULM-based microvascular information have already shown promise for pathological applications, including stroke detection [14] and longitudinal monitoring [15]. Volumetric ULM recovers the vascular structure and flow information from the same acquisition by localizing and tracking intravenously injected microbubbles[16–19]. Vascular features, such as vessel density, flow velocity, vessel diameter, tortuosity, and local flow rate, can then be extracted [20]. In disease or intervention studies, multiple aspects of the vasculature may change. Jointly analyzing these spatially distributed measurements produces high-dimensional data that are difficult to interpret. Testing each feature separately across brain locations increases the multiple-comparison burden and may obscure coordinated vascular changes when individual effects are modest. On the other hand, interpreting significant features one at a time does not reveal how structural and flow-related measures are jointly organized.

High-dimensional biological measurements often contain coherent low-dimensional organization. This can be unveiled by use of dimensionality reduction techniques. One direct approach is to summarize this organization by dividing the brain into discrete regions using clustering algorithms, such as k-means and spectral clustering [21]. Many brain properties, however, vary continuously across space, such as gene expression and myelin content [22]. Hard boundaries imposed on fundamentally continuous organization can obscure the underlying structure. Gradient methods offer an alternative. They describe the data using a few dominant axes along which organization changes smoothly. This can be achieved with linear methods such as PCA, or with nonlinear methods such as affinity-based manifold learning. Nonlinear methods have been widely adopted in recent years, as they capture indirect and direct relationships between brain regions and show higher sensitivity to locally, nonlinearly-embedded structure than linear methods [23]. These dominant axes are commonly termed gradients, emphasizing their smooth and graded nature, though, formally, in the nonlinear case, each axis is a coordinate on a lower-dimensional manifold [24,25].

Gradient organization has been described across a wide range of brain properties in humans, including dominant axes of cortical gray matter [26] and white matter [27] functional organization, cortical microstructure [28] and myeloarchitecture [29], and cerebellar structural and functional organization [30,31]. Similar gradient organization has also been reported in rodents, including brain functional organization [32], gene expression [33,34], and cytoarchitecture [35].

Gradient-like organization of cerebrovasculature remains underexplored. Most cerebrovascular studies have focused on structural vascular atlases, local hemodynamic measurements, or global perfusion. In human infancy, the developmental trajectory of cerebral blood flow has been reported to emerge along a similar limbic-sensorimotor-association hierarchy that organizes cortical functional and metabolic specialization [36]. More recently, gradient organization of adult human cerebral perfusion was reported at mesoscale, revealing a principal axis dominated by upstream vascular territory [37]. Whether such organization extends to the microvascular scale, and whether local vascular structure and flow dynamics jointly define coherent gradients in vivo, remains unclear.

Here, we characterized high-dimensional local vascular phenotypes derived from volumetric ULM across the mouse brain and tested whether they exhibited reproducible low-dimensional organization. We constructed a group-consensus vascular phenotype graph from ULM-derived local vascular structure and flow dynamic measures, where brain regions with similar vascular profiles were more strongly connected. Diffusion embedding was applied to this graph to identify its dominant gradients, that is, coordinates on a lower-dimensional manifold. We then mapped the resulting gradients across the brain, summarized their positions across networks, and examined the vascular features associated with each axis. We show that in vivo cerebrovascular structure and flow are organized along low-dimensional gradients, providing a reference and framework for investigating coordinated vascular reorganization in future longitudinal, interventional and disease-model studies.

## 02 Results

### Volumetric ULM provides a high-dimensional vascular phenotype

To define the vascular phenotype space for group-level analysis, volumetric ultrasound localization microscopy was performed in each mouse, and the reconstructed datasets were rigidly aligned to a common space. The ULM data provided a high-dimensional local vascular phenotype. From the same acquisition, at each spatial location, this phenotype comprised distributional summaries of local vascular density (bubble count), tortuosity (sum-of-angles metric), and blood flow speed, acceleration (acceleration and deceleration), and directional complexity (local angular deviation, turning angle). Supplementary Figure S2 shows the subject-wise feature distributions.

Representative ULM maps showed complementary aspects of the vascular signal obtained from the same acquisition (**Figure 1**a–c). The directional density map resolved vascular structures according to the direction of tracked microbubble flow (Figure 1a). The speed map and flow direction map showed spatial variation in the magnitude and orientation of local flow velocity, respectively (Figure 1b and c). These local multi-feature profiles were then used to test whether the vascular phenotype data contained a lower-dimensional organization shared across mice.

**Figure 1.**
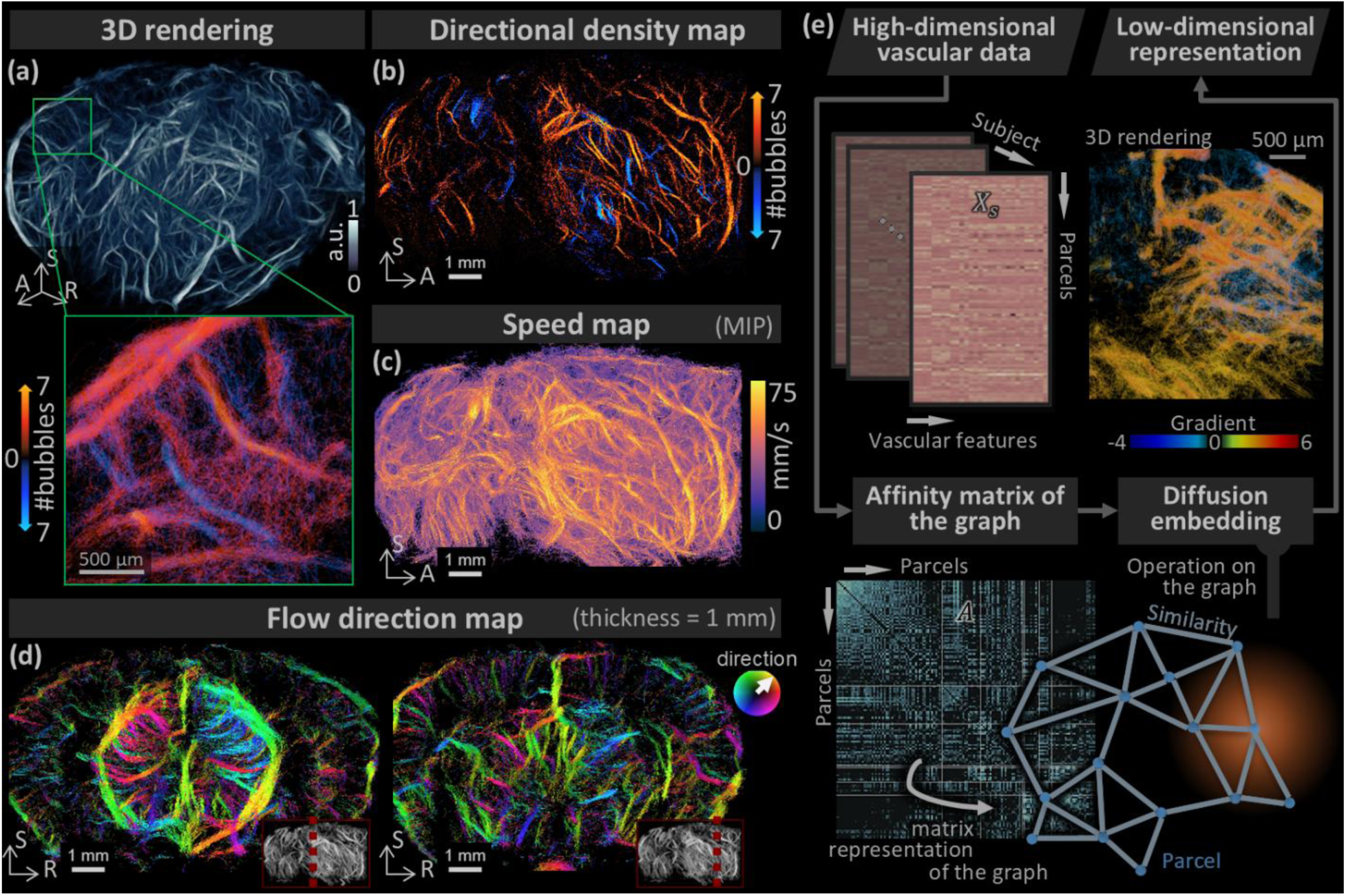
From volumetric ULM vascular phenotype data to low-dimensional vascular gradients via diffusion embedding. Volumetric rendering of the ULM density map (a), volume maximum-intensity projections (MIP) of the directional density map (b) and speed map (c), and 1 mm slabs of flow direction map (d), from a representative mouse. (e) For each subject, parcel-to-parcel vascular similarity is computed from their parcel-by-feature matrix ***X_s_***. These subject-specific similarity matrices are combined across subjects into a group-consensus affinity matrix ***A***. It defines a graph where nodes are parcels and edge weights reflect pairwise vascular similarity. Diffusion embedding is applied to this graph to extract the gradients **Ψ*_k_***: a ranked, low-dimensional set of axes on a manifold where proximity reflects vascular similarity. In panel (b) – (d), all scale bars indicate 1 mm. A: anterior; R: right; S: superior.

### Group-consensus embedding identifies two coherent low-dimensional vascular phenotype gradients

To test whether the high-dimensional ULM-derived vascular phenotype contained a lower-dimensional organization shared across mice, we constructed a group-consensus vascular phenotype graph and applied a nonlinear dimensionality reduction method, diffusion embedding (Figure 1d). A group-level ULM density map was first used to generate a spatially compact, vessel-informed parcellation of the brain volume. This defined a common set of 150 brain segmentations, termed parcels, for summarizing local vascular phenotypes across mice (Figure 1d). Supplementary Figure S3 and S4(a) show the group-level ULM density map and the parcellation result, respectively.

For each mouse, the high-dimensional phenotype profile was summarized within each parcel, and parcel-to-parcel similarity was calculated from the resulting parcel-by-feature matrix. Subject-specific similarity matrices were then combined to generate a group-consensus affinity matrix (Supplementary Figure S4(b)), where parcels with similar vascular phenotypes were more strongly connected. Diffusion embedding of this affinity matrix identified a low-dimensional arrangement of parcels according to their multivariate vascular phenotype similarity.

The eigenvalue spectrum indicated that the first two embedding dimensions captured the dominant structure (Figure 1d and Supplementary Figure S4(b)). We next tested whether these dimensions were robust across individuals and across spatial scales. In leave-one-mouse-out analyses, the leading gradients remained highly correlated with those derived from the full group-consensus matrix, with correlations greater than 0.928 for the first three gradients, and correlations greater than 0.856 for the top five gradients (Supplementary Figure S5). The first two gradients were also preserved across parcellation, with correlations greater than 0.809 for the principal gradient and 0.769 for the first two gradients ranging from 125 to 200 parcels (Supplementary Figure S5). Thus, the two leading vascular phenotype gradients were reproducible across mice and across parcellation scales. We therefore focused on these gradients for subsequent spatial and phenotypic interpretation.

### The two vascular phenotype gradients have distinct spatial distributions and regional positions

To characterize the spatial organization of the two leading vascular phenotype gradients, parcel scores were mapped from the embedding to the common imaging space (**Figure 2**). The principal gradient was broadly expressed across cortical and subcortical territories, whereas the secondary gradient showed a more prominent separation between isocortical and deep subcortical regions (Figure 2a, b).

**Figure 2.**
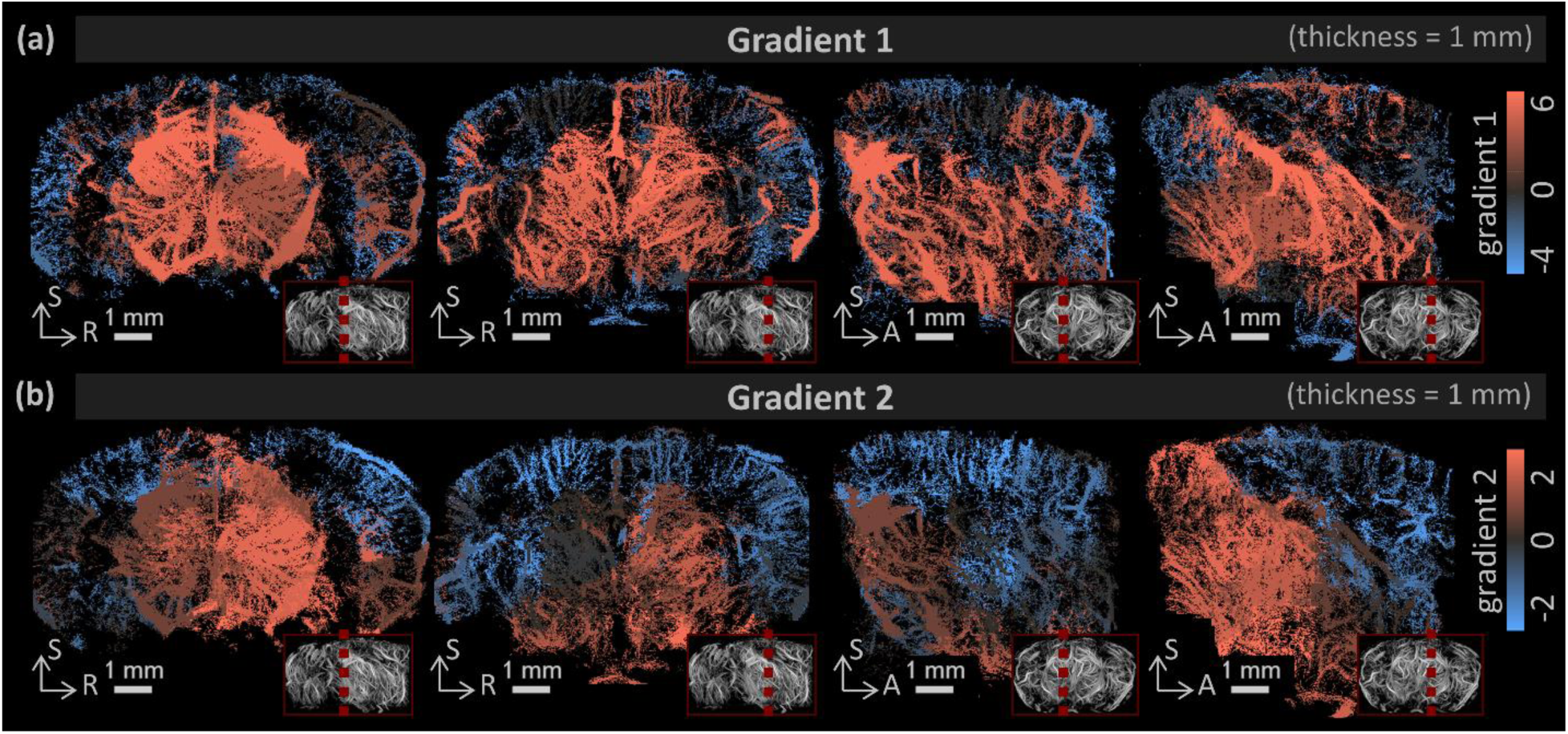
The principal (a) and secondary (b) vascular phenotype gradients show distinct spatial distributions across the brain volume. Figures are shown as color-coded 1 mm slabs. The thumbnail in the red box indicates the slab location. All scale bars indicate 1 mm. A: anterior; R: right; S: superior.

We next summarized the gradients within major brain regions defined by the Alan atlas, including isocortex, hippocampus, striatum, thalamus, hypothalamus, and midbrain (**Figure 3**). In the two-gradient embedding space, isocortex, hippocampus, and striatum occupied positions toward one end of the joint distribution, whereas thalamic, hypothalamic, and midbrain parcels were shifted toward the other end (Figure 3a). This pattern provided a broad separation between telencephalic and deeper diencephalic and mesencephalic territories, while retaining substantial variation within individual regions.

**Figure 3.**
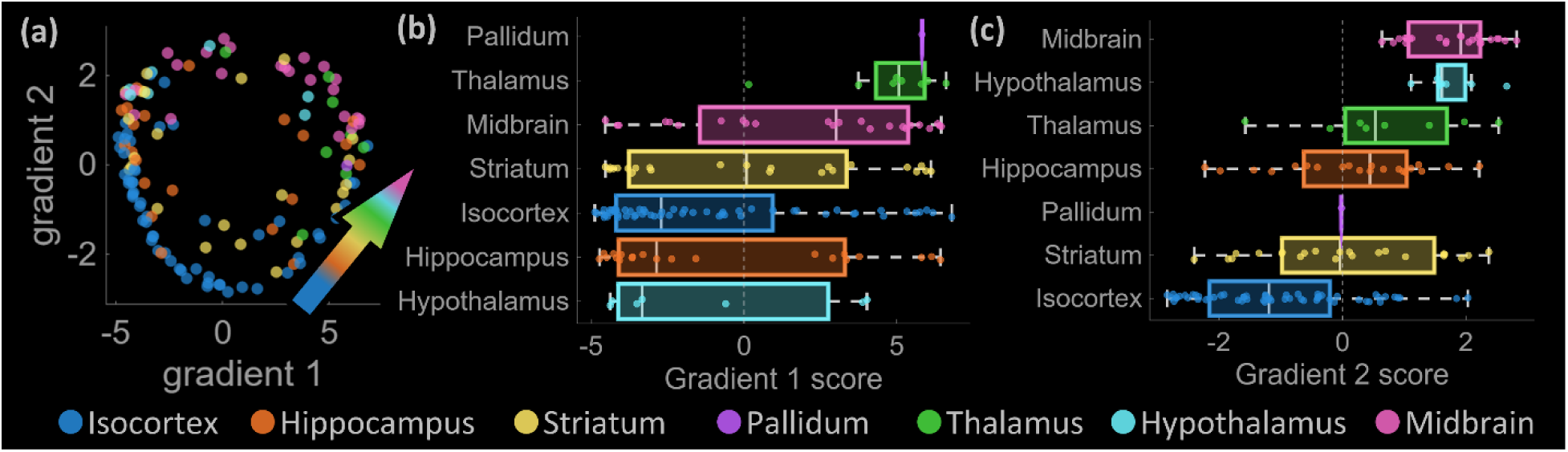
Vascular phenotypical gradients show a cortical-to-deep-subcortical ordering, from telencephalic (isocortex, hippocampus) to diencephalic (thalamus, hypothalamus) to mesencephalic (midbrain). (a) Regional separation in the space defined by the top two vascular phenotypical gradient. (b) Region-wise distribution of gradient 1 scores, ordered by regional median. (c) Region-wise distribution of gradient 2 scores, ordered by regional median. In all panels, each data point is the gradient score of a parcel, and all data points are colored by brain region

Along the principal gradient, regional score distributions overlapped broadly (Figure 3b). In contrast, the secondary gradient showed clearer network separation. Hypothalamic and midbrain regions anchored one end of the axis, whereas isocortical regions were positioned toward the other (Figure 3c). The hippocampus, striatum, and thalamus occupied intermediate and partially overlapping positions along the secondary gradient.

### The principal gradient defines a vascular structural-regime axis

To characterize the vascular phenotype underlying the principal gradient, we quantified the association between Gradient 1 scores and the ULM-derived local phenotype features in each mouse (**Figure 4**a). Gradient 1 scores were positively associated with local vascular density features (local density) and negatively associated with structural complexity features (turning angle, angular deviation). Flow dynamic features, including acceleration- and deceleration-related features, also showed weak negative associations with the gradient. These associations were consistent across subjects.

**Figure 4.**
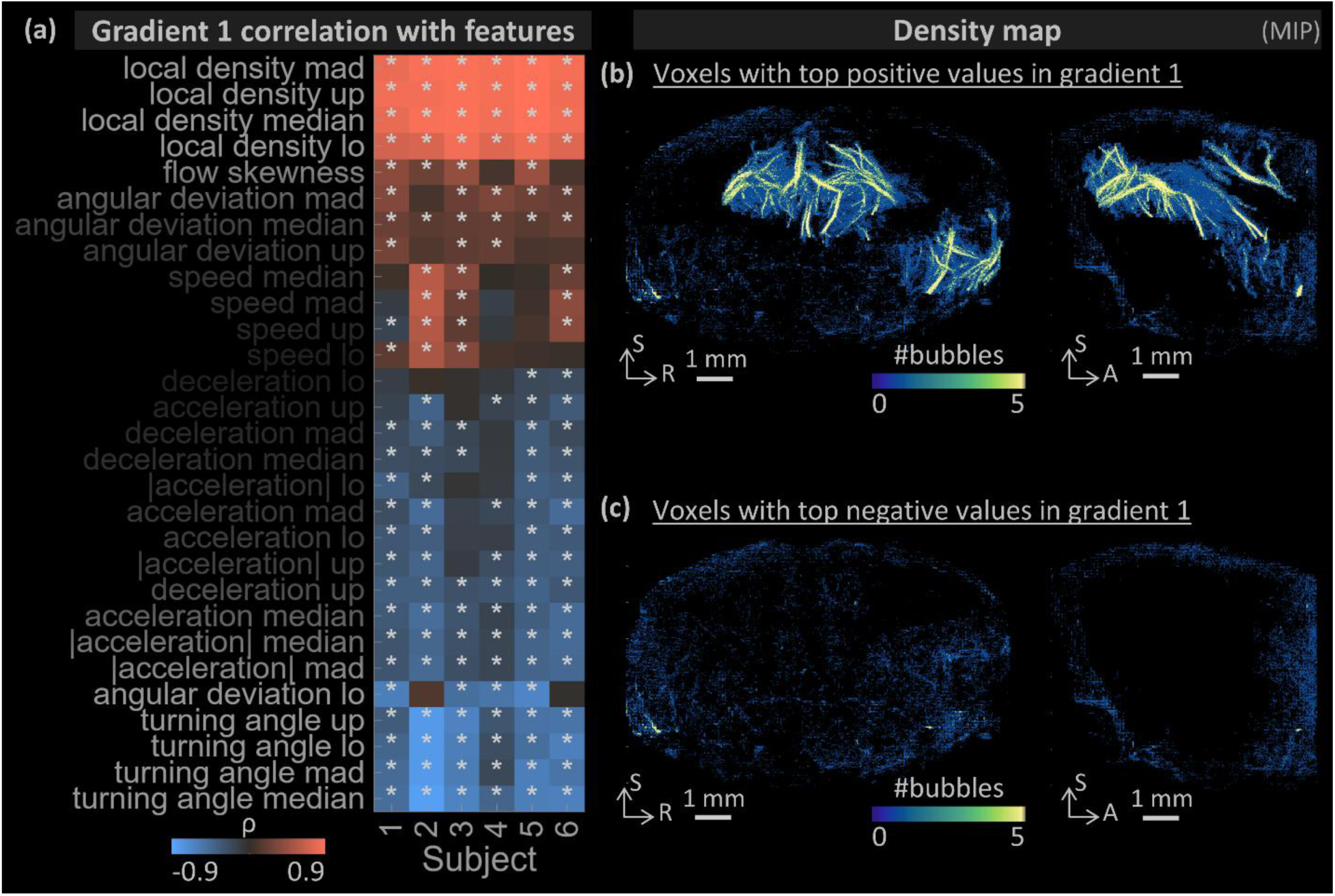
Principal gradient defines a vascular structure-related axis. (a) Subject-wise correlations between gradient 1 and parcel-level vascular features, ordered by mean correlation across subjects. (b) Density map of the voxels with top 5% positive gradient 1 values. (c) Density map of the voxels with top 5% negative gradient 1 values. In panel a, white star indicate the FDR-corrected p-vlaues smaller than 0.05. lo: 5th percentile; up: 95th percentile; mad: median absolute deviation.

To examine the spatial correspondence between Gradient 1 and local vessel structure, we visualized brain regions at opposing ends of Gradient 1 on the ULM density map in the common imaging space. Brain regions with the highest positive Gradient 1 scores were preferentially located within major vascular structures visible in the ULM density map (Figure 4b). In contrast, brain regions with the most negative Gradient 1 scores were distributed across lower-density vascular territories and showed a broadly distributed spatial pattern (Figure 4c).

Thus, these findings indicate that the principal gradient defines a vascular structural regime. One end of the principal gradient was associated with higher local vascular density and more directionally coherent flow, whereas the other end of was associated with lower density and more complex vascular phenotypes.

### The secondary gradient defines a flow-dynamics-regime axis

To characterize the vascular phenotype underlying the secondary gradient, we quantified the association between Gradient 2 scores and the ULM-derived local phenotype features in each mouse (**Figure 5**a). Gradient 2 scores were positively associated with flow dynamics features (speed, deceleration, acceleration) and negatively associated with flow complexity features (turning angle). These associations were generally consistent across subjects, although they were less consistent than those observed for Gradient 1, particularly at the negative end of Gradient 2.

**Figure 5.**
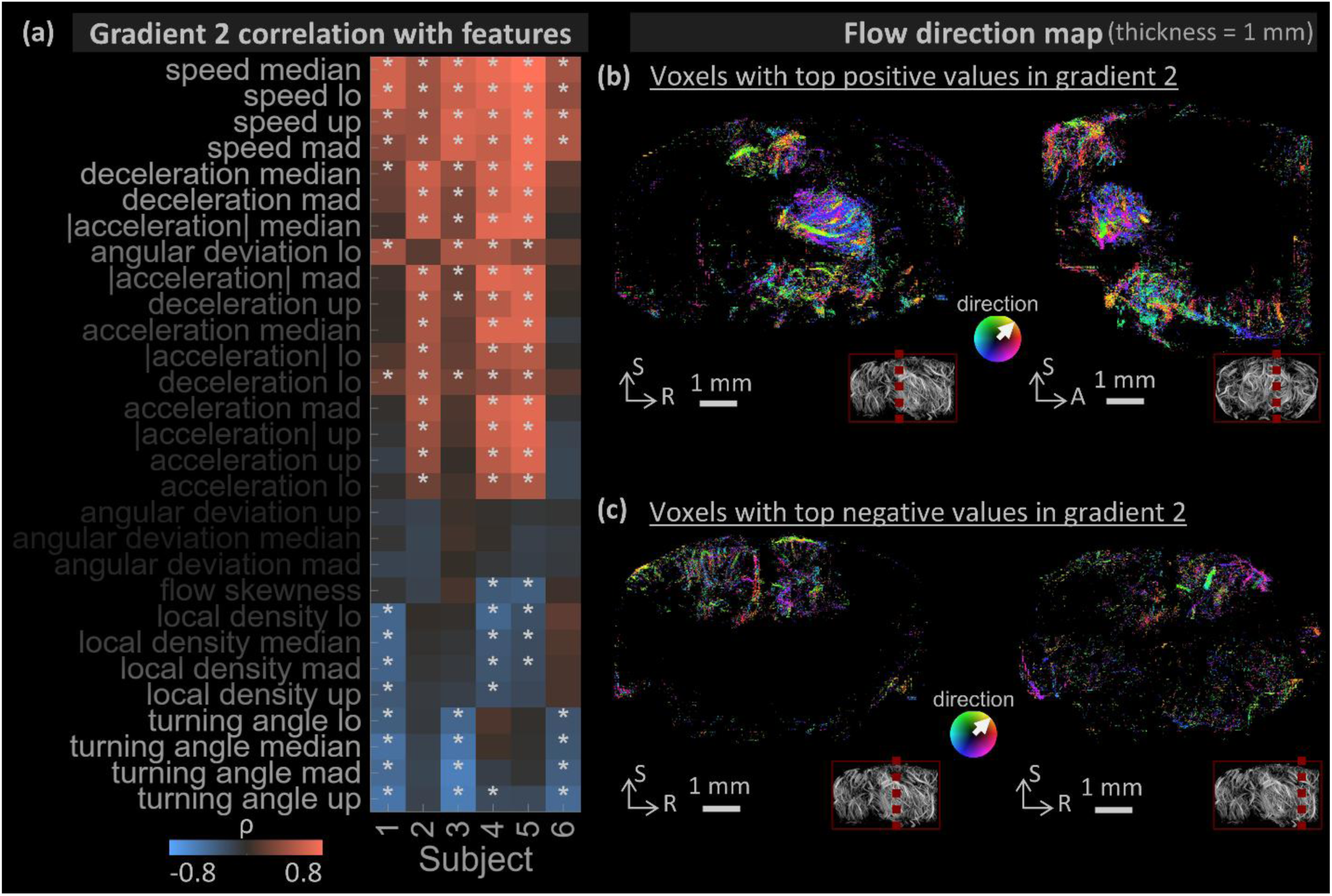
Secondary gradient defines a flow dynamics-related axis. (a) Subject-wise correlations between gradient 2 and parcel-level vascular features, ordered by mean correlation across subjects. (b) Flow direction map of the voxels with top 5% positive gradient 2 values. (c) Flow direction map of the voxels with top 5% negative gradient 2 value. In panel a, white star indicate the FRD-corrected p-vlaues smaller than 0.05. In panels b and c, the thumbnail in the red box indicates the slab location. All scale bars indicate 1 mm. A: anterior; R: right; S: superior. lo: 5th percentile; up: 95th percentile; mad: median absolute deviation.

To examine the spatial correspondence between Gradient 2 and local flow organization, we visualized brain regions at opposing ends of Gradient 2 on the ULM velocity direction map in the common imaging space. Brain regions with the highest positive Gradient 2 scores were located primarily in subcortical areas, particularly the midbrain and hypothalamus (Figure 5b). In contrast, brain regions with the most negative Gradient 2 scores were located predominantly in isocortical areas (Figure 5c).

Thus, these findings indicate that the secondary gradient defines a flow-dynamics regime. One end of the secondary gradient was associated with faster and less complex blood flow in the subcortex, whereas the opposite end was associated with slower and more complex flow in the isocortex.

## 03 Discussion

In this study, we showed that high-dimensional vascular phenotypes derived from volumetric ULM can be organized into reproducible low-dimensional gradients across the mouse brain. Using group-consensus diffusion embedding, we identified two dominant axes, termed gradients, that were preserved across individual and spatial scales. The principal gradient was associated with local vascular structure, spanning from higher density to more tortuous structure, whereas the secondary gradient was associated with local flow dynamics, spanning from higher flow to greater trajectory complexity. These gradients organized continuously across the brain volume, and showed distinct spatial distributions and network-level positions. Together, these findings extend volumetric ULM from regional visualization and single-feature quantification toward a multivariate representation of in vivo vascular phenotyping.

Volumetric ULM images contain high-dimensional vascular structure and flow dynamics data that are difficult to interpret, and feature-wise analyses may obscure coordinated variation across brain regions. Here, we present a framework for extracting and interpreting low-dimensional vascular organization from high-dimensional volumetric ULM data. By using diffusion embedding, we reduced these multi-feature measurements to two continuous axes that were reproducibly recovered across individuals and spatial scales. Similar gradient-based approaches in MRI been used to characterize developmental changes in large-scale brain organization [27,55] and age-related reorganization of morphometric brain networks [56]. Applied to volumetric ULM, the framework developed here could provide a reference phenotype space for testing how cerebrovascular disease, aging, or ultrasound-based interventions alter the joint organization of cerebrovascular structure and flow, rather than affecting each vascular feature separately within each distinct network.

The principal gradient revealed an axis that was associated with larger and more structured vessels at one end, and with a broadly distributed capillary network and greater trajectory complexity at the other. This pattern resonates with known regional heterogeneity in the mouse cerebrovasculature, where regional differences lie primarily in capillaries and large vessels [7]. It also agrees with the cerebrovasculature reconstructed using micro-optical sectioning tomography by Ji and colleagues [8]. They identified vessel length density as a dominant vascular structural parameter, linked to the local metabolic demand [8]. A conceptually related axis has been described with ASL in humans, where a perfusion gradient differentiated core arterial supply territories from border zones [37]. Because the brain contains a distributed capillary network supplied by larger upstream vessels, it is not surprising that both poles of the principal gradient were distributed across multiple brain networks. Even so, the regional summary showed that the hypothalamus and hippocampus weighed strongly toward the capillary-dominated end, whereas the thalamus weighed toward the larger-vessel end. This agrees with post-mortem data showing that the thalamus and primary somatosensory cortex have high capillary density, whereas the hippocampus has lower capillary density and the hypothalamus contains comparatively fewer medium- to large-sized vessels [7]. This is also consistent with the findings in awake mice that the hippocampus has a lower capillary density, which is associated with lower red blood cell speed and flux and lower resting blood oxygenation compared to the neocortex [57]. Thus, the principal gradient captured a vessel structure axis. It described a continuum from regions dominated by larger, directionally organized vessels to regions with more capillary-like and tortuous vascular patterns. Given the established relation between regional vascular architecture and metabolic support, this axis may also relate to the spatial organization of local metabolic demand across the brain, although this possibility requires independent metabolic validation.

The secondary gradient revealed an axis that was associated with higher flow speed, highlighting the hypothalamus and midbrain regions at one end, and with stronger anisotropic flow, highlighting the isocortex at the other. This agrees with previous reports of regional variation in capillary length, branching, and orientation [8]. Capillary branches are shorter in the striatum and hypothalamus and longer in the somatosensory cortex and thalamus. Capillary anisotropy is low in the inferior colliculus. In the cortex, the capillaries are preferentially aligned parallel to cortical layers and therefore perpendicular to the penetrating cortical vessels. This pattern is depth-dependent, with strong perpendicular orientation in superficial layers and weaker orientation in deep layers [8]. These structural observations may offer one possible mechanistic basis for the observed regional separation in this flow dynamic-regime axis. The cortical pole, characterized by larger turning angles indicating a strong anisotropic flow, is supported by a highly organized, layer-dependent capillary architecture. The subcortical pole, characterized by higher flow speed, corresponds to regions with shorter, more isotropic capillary networks that may offer lower vascular resistance. Furthermore, the midbrain and hypothalamus are supplied by the posterior cerebral and anterior choroidal arteries, whereas the isocortex is supplied by the anterior and middle cerebral arteries [7]. This separation is conceptually related to the principal perfusion gradient identified in humans, which separates the posterior and the anterior circulation territories [37]. Thus, the secondary gradient captured a flow dynamics axis. It differentiates subcortical vascular phenotype from cortical vascular phenotype. Its biological basis remains to be established, but may involve regional differences in capillary architecture at the microvascular level and the anterior-posterior arterial territory division at the macrovascular level.

Several limitations are worth considering. The analysis was based on female C57BL/6J mice imaged under isoflurane anesthesia. The generalizability of the derived gradients requires further testing in larger cohorts, across sexes, ages, strains, and physiological states. Uneven microbubble detection and incomplete coverage across the brain also limited the analysis to regions reliably sampled across all mice. Consequently, the present gradients describe the imaged cerebrum and brainstem rather than the entire brain. Rigid registration provided a conservative common space but did not fully account for inter-animal anatomical differences. This is a compromise, as the nonrigid warping could have distorted vascular geometry and track-derived features. Nevertheless, the leading gradients remained stable after exclusion of individual mice and across parcellation scales, indicating that the observed low-dimensional organization was not random. In this healthy wild-type cohort, these results provide a reference representation of cerebrovascular structure and flow. Future studies could test whether disease produces a shift, compression, expansion, or spatial reorganization of the vascular phenotype gradients. Such analyses may help distinguish coordinated brain-wide vascular reorganization from localized changes in individual ULM measures.

In conclusion, volumetric ULM combined with group-consensus diffusion embedding revealed two reproducible gradients of vascular phenotype in the mouse brain. These gradients provide a compact representation of in vivo cerebrovascular structure and flow and establish a reference phenotype for future studies of vascular change across aging, disease, and intervention.

## 04 Methods

### Animals

Ten wild-type mice (Envigo, C57BL/6J) were used in this study. Four were used for protocol optimization. Six were included in the formal analysis. They arrived one week before the experiment, and were housed under controlled conditions. All procedures in this paper were approved by the UK Home Office and Imperial College London’s Welfare and Ethical Review Body (PPL: PBDB8AA12). Animal experiments were conducted following the UK Animals (Scientific Procedures) Act 1986 and ARRIVE guidelines [38].

Mice were anesthetized with 5% isoflurane in an induction chamber. Fur over the imaging window, extending from bregma to the neck, was removed by shaving followed by depilatory cream (Nair Sensitive). Each animal was placed on a heating mat and secured in a stereotaxic frame with ear bars to minimize head motion. Isoflurane was maintained at 1-2% through a face mask during imaging. A water bowl was positioned over the imaging window. Homemade microbubbles were prepared as described previously [39], and infused through a 30-gauge tail-vein catheter using a custom-made rocking syringe pump at 0.3 μl/min/g. The total injection volume did not exceed 250 μl for a 30 g mouse. The total duration of each imaging session did not exceed 1 hour. After imaging, mice received oxygen and recovered in a warmed enclosure until normal activity resumed before returning to their home cages.

### Imaging Acquisition

A Verasonics Vantage 256 system (Verasonics Inc., Kirkland, WA, USA) and a multiplexed 8 MHz matrix probe of 1024 elements (Vermon, Tours, France) were used for all acquisitions. Plane waves were transmitted at the center frequency without steering angles using the full aperture and were received sequentially by four sub-apertures. The transmit voltage was set at 30 V to compensate for the skull attenuation to achieve an MI of 0.05. For each acquisition, 400 image blocks were recorded. Each image block consisted of 400 volumes over 1 s duration. The full acquisition lasted 7 min. This frame rate, 400 Hz, was chosen as the highest rate that allowed continuous data streaming and saving with the available Verasonics system, host computer, and solid-state drive hardware.

### Image reconstruction

RF data was beamformed using the delay-and-sum with an isotropic voxel size of one quarter of the pitch, that is, 75 μm. Each acquisition was reconstructed as consecutive blocks of 400 volume frames. The beamformed complex data contained tissue, microbubble, and noise signals and were processed separately for each block. For clutter filtering, beamformed complex data were reshaped into a Casorati matrix whose dimensions corresponded to the number of voxels by the number of frames. The Casorati matrix was then decomposed using SVD into left singular vectors, singular values, and right singular vectors. The clutter threshold was selected as the elbow point in the plot of singular-value magnitudes using the publicly available MATLAB function kneept [40]. The noise threshold was determined based on the frequency curve of the singular vectors. Typically, the first 10 components were removed to obtain bubble signals, and the last 10 components were used for noise estimation. The bubble signal was reconstructed from the remaining components.

To reduce spatially inhomogeneous noise, the reconstructed bubble signal was further equalized using a noise field. The spatial noise field was estimated as the temporal median magnitude of the reconstructed highest-order SVD components and spatially smoothed with a Gaussian filter. The equalization map was computed as the ratio between a reference noise level, defined here as the median of the noise field, and the local noise estimate. This map was then applied to the reconstructed bubble signal to compensate for spatially varying noise and produce a more uniform background for subsequent localization and tracking. Supplementary Figure S1 shows the results with and without the noise equalization step.

### ULM Image Reconstruction

De-cluttered frames were thresholded at 0.05 of the maximum image intensity. For each dataset, a representative point spread function (PSF) was generated by averaging five manually selected isolated bubbles within this dataset. Microbubbles were localized with a cross-correlation-based method and tracked using a Kalman filter and graph-based algorithm [41,42]. Tracks shorter than 15 frames were discarded. Bubble tracks were accumulated over all 160, 000 frames to reconstruct the density map. Parameters were empirically chosen for a balance between the preserved bubble tracks and false tracks.

### Atlas Registration

Brain region segmentation was based on the Allen Mouse Brain Common Coordinate Framework version 3 [43]. To warp the atlas segmentation into the imaging space of this study, a vascular reference dataset available in Allen space [44] was registered to a vascular reference volume of this study. The reference volume was derived from a temporal maximum-intensity projection of the CEUS data from one representative mouse in this study. The registration was performed in 3D Slicer (https://www.slicer.org/ [45]) using ANTS (Rigid + Affine, default parameter settings [46]). The resulting transformation was then applied to the Allen brain region segmentation to warp the atlas labels into the imaging space of the present study.

### ULM Registration

A two-step registration pipeline was used to align the volumetric ulm data across mice. First, each dataset was manually aligned to correct gross differences in orientation. Second, rigid landmark-based registration was performed using anatomically corresponding landmarks.

A set of 20 reference landmarks was defined in the reference mouse. Corresponding landmarks were then selected in each of the other mice only when they were clearly identifiable on visual inspection; otherwise, the landmark was omitted. For each registration, at least 5 landmark pairs were required.

Rigid transformation was estimated using the MATLAB function fitgeotform3d [47]. The final transformation combined the manual alignment and the rigid landmark-based transform, and was applied in a single resampling step using imwarp with linear interpolation and the reference image grid as the output space.

Only rigid registration was used in the main analysis. This choice was made for two reasons. First, anatomical mapping was incomplete across datasets, which limited the reliability of more complex transformations. Second, non-rigid registration would deform vessel geometry and could therefore alter the quantification of bubble tracks.

### ULM-based metrics extraction

Vascular features were computed for each track, and where applicable, for each track section.

The speed was computed for each track section *i* as *v_i_* = Δ*x_i_* × FR, where the Δ*x* is the displacement length of the given track section *i* and FR is the frame rate. The acceleration was defined as the first derivative of speed between consecutive track sections *a_i_* = Δ*v_i_* × FR. In the results section, the absolute acceleration |*a_i_*|, the magnitude of positive acceleration |*a_ven_*|, and the magnitude of negative acceleration |*a_art_*| were analyzed separately. The tortuosity was quantified by the soam, defined as 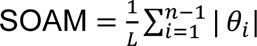., where *L* is the length of the track trajectory, *n* is the number of track sections, and *θ* is the angle between consecutive track sections [48]. The angular deviation was defined as the angle between an individual track direction and the principal flow direction of the voxel through which that track passed. The principal flow direction was defined as the first component obtained by applying pca to all flow vectors associated with that voxel [49]. The turning angle was defined as the angle between two consecutive track sections.

### Group-consensus parcellation

To characterize the dominant low-dimensional organization of the high-dimensional vascular phenotype data across individuals, a group-consensus, vessel-aware mesoscale parcellation was first generated from group-level vessel density maps. This parcellation defined a common set of nodes across subjects, that is, the nodes of a group-level graph, where each edge weight reflects the vascular phenotype similarity between two parcels. Diffusion embedding was applied to this graph to uncover its dominant axes of organization, termed gradients (see for known gradient organizations in the brain).

A group-level ULM density map was computed by averaging the density maps across subjects ((a)). This group-level ULM density map was used as a soft guide in the parcellation so that parcel boundaries were influenced by vascular support while remaining primarily spatially compact. Specifically, voxels were represented by their spatial coordinates and their bubble density, and partitioned into a predefined number of clusters using k-means clustering [50]. The number of clusters was tested from 50 to 200 in increments of 25. To ensure the connectivity of the clusters, for each parcel, only the largest connected component was retained under the original label, while smaller disconnected fragments were reassigned to neighboring parcels. (b) shows the parcellation results, and (a) shows the size of the 150 parcels organized by brain regions.

### Affinity matrix construction

For each subject, a parcel-by-feature matrix was constructed by summarizing multiple local vascular features within each parcel, including the local median, mad, and 5th and 95th percentiles from density, speed, angular deviation, turning angle, acceleration, and deceleration, and histogram skewness of the first derivative of speed.

Parcel-to-parcel similarity matrices were then computed separately for each subject from their parcel feature profiles, as *R_s_* = corr(*X_s_*), where *X_s_* is subject *s*’s z-scoredd parcel-by-feature matrix, and corr(⋅) denotes the Pearson correlation between each parcel pair across all feature profiles. Subject-specific similarity matrices were then combined into a group-level similarity matrix (*R̅*) using Fisher z-transformation, averaging in z-space, and back-transformation to correlation space, as 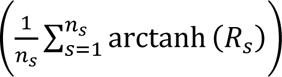, where *n_s_* is the number of subjects. Fisher z-averaging was used because the distribution of Pearson correlation coefficients is skewed and this operation standardizes the variance, making it a less biased choice than direct averaging [51]. The resulting group similarity matrix was then converted to a non-negative affinity matrix, *A* = max(*R̅*, 0) ((b)), as diffusion embedding requires non-negative edge weights to define a valid Markov transition matrix. Here, negative correlations were treated as absence of shared vascular phenotype.

### Diffusion embedding

This affinity matrix was then used for diffusion embedding using BrainSpace [52]. Diffusion embedding is an unsupervised, affinity-based manifold learning algorithm that uncovers the intrinsic geometry of a graph [24,25]. The name reflects two ideas. Diffusion, because the algorithm models heat diffusion on a graph via a random walk (Markov chain), where a walker moves between nodes with probability proportional to their affinity. Embedding, because the geometry of this diffusion process is used to place each node into a low-dimensional coordinate space that preserves the diffusion distances between them. Here, this gives a set of coordinates, that is, the gradients, describing the manifold geometry that best reflects the global structure of vascular similarity across brain regions. Regions with similar vascular profiles are positioned close together, and dissimilar regions are positioned far apart. Detailed mathematical derivation of diffusion embedding can be found in two Coifman papers [24,25].

Briefly, an anisotropic kernel, *A*^(*α*)^, was computed to correct for uneven nodal degree, so that the resulting connection strengths reflect the intrinsic geometry of the graph rather than how broadly connected a node is to the rest of the graph. This anisotropic kernel was computed as *A*^(*α*)^ = *D*^−*α*^ *A D*^−*α*^, where *D* is the diagonal degree matrix of *A*, with diagonal entries *D_ii_* equal to the degree of node *i*, that is, ∑*_j_ A_ij_*, and *α* ∈ [0,1] controls the strength of this correction. *α* = 0 means no correction, equivalent to the standard graph-Laplacian normalization, while *α* = 1 removes degree influence entirely, considering only the geometry of the data, equivalent to the Laplace-Beltrami operator. Here, the default value, *α* = 0.5 (backward Fokker–Planck operator), was used, consistent with the gradient literature in neuroscience [26,53]. It partially removes the influence of nodal degree on the graph, balancing between the raw connection structure and the intrinsic geometry of the data.

The diffusion operator was then computed as *M* = *D*^(*α*)−1^ *A*^(*α*)^, where *D*^(*α*)^ is the diagonal degree matrix of *A*^(^*^α^*^)^, with diagonal entries *D*^(^*^α^*^)^*_ii_* equal to the degree of node *i*, that is, 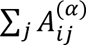. *M* defines the Markov chain, where *M_ij_* is the probability of a random walk stepping from node *i* to node *j* in a single step. The gradients were derived through eigendecomposition of *M*: *Mφ_k_* = *λ_k_φ_k_*. The *k*-th gradient was then computed as 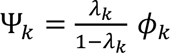, where *φ_k_* and *λ_k_* are the *k*-th eigenvector and eigenvalue of *M* (ranked *λ*_1_ ≥ *λ*_2_ ≥ ⋯). This *λ_k_*/(1 − *λ_k_*) scaling is controlled by the diffusion_time parameter in BrainSpace. diffusion_time = 0 was used, consistent with the gradient literature in neuroscience [26,53]. This setting performs an automated multi-scale estimation by summing the diffusion process analytically over all step counts rather than fixing one, making it more robust to noise and data distribution than fixing a single diffusion time.

## Statistical Analysis

To assess the reproducibility of the vascular phenotype gradients across individuals, leave-one-subject-out analyses were performed. Comparing the gradients derived from the group-level affinity matrix to gradients derived from affinity matrices with leave-one-subject-out, the top five gradients remain stable ((a), *ρ* > 0.928 for the top 3 gradients, *ρ* > 0.856 for the top 5 gradients).

To assess sensitivity to the spatial scale used to summarize the ULM phenotype, the full analysis pipeline was repeated using group-consensus parcellations ranging from 50 to 200 parcels with 25 increments. Gradients derived from each parcellation scheme were compared with those from the 150 parcels using Pearson correlation. The 150-region solution was selected for the main analysis because the first two gradients were preserved across nearby parcellation schemes while retaining sufficient spatial resolution for anatomical interpretation.

To aid interpretation, parcel gradient scores were summarized into larger brain regions, namely isocortex, hippocampus, striatum, thalamus, hypothalamus, and midbrain, using the majority-overlap parcel-to-region assignment. To characterize the vascular phenotype associated with each gradient, Spearman correlations were calculated between gradient scores and each ULM-derived local phenotype feature across brain regions, for each mouse. Statistical significance was assessed using two-sided tests, with false-discovery-rate correction applied across all feature associations and mice within each gradient using publicly available MATLAB function fdr_bh [54].

All the data processing and visualization in this study were conducted in MATLAB (R2023b, MathWorks, Natick, MA, USA).

## Acknowledgements

We would like to thank all the animals and animal technicians for making this study possible. This project is supported by the Chinese Government Scholarship (CSC). Additional funding is provided by the Advanced Research and Invention Agency (ARIA), UKRI/Wellcome grant EP/W024020/1 and Wellcome 338562/Z/25/Z.

## CRediT authorship contribution statement

J.Z. conceptualized the project, acquired data, processed and analyzed the data, and drafted and revised the manuscript. S.K. acquired data and revised the manuscript. K.R. were consulted regarding data acquisition and imaging reconstruction. J.Y. provided key code for ultrasound localization microscopy. B.W. provided key code for data acquisition and imaging reconstruction. S.V.M. were consulted regarding data acquisition. S.R.S. and M.-X.T. oversaw its execution, provided insights, and edited the manuscript. All authors edited and approved the final version of the manuscript.

## Declaration of generative AI use

During the preparation of this work the authors used ChatGPT (OpenAI) and Claude (Anthropic) to help develop code and polish text. After using this tool, the authors reviewed and edited the content as needed and take full responsibility for the content of the published article.

## Data and Code Availability

Raw data and code reported in this study are available upon reasonable request to the corresponding author.

