## Supplementary Figure for "Cerebrovascular Structure–Flow Gradients Revealed by Volumetric Ultrasound Localization Microscopy"

### Supplementary figures

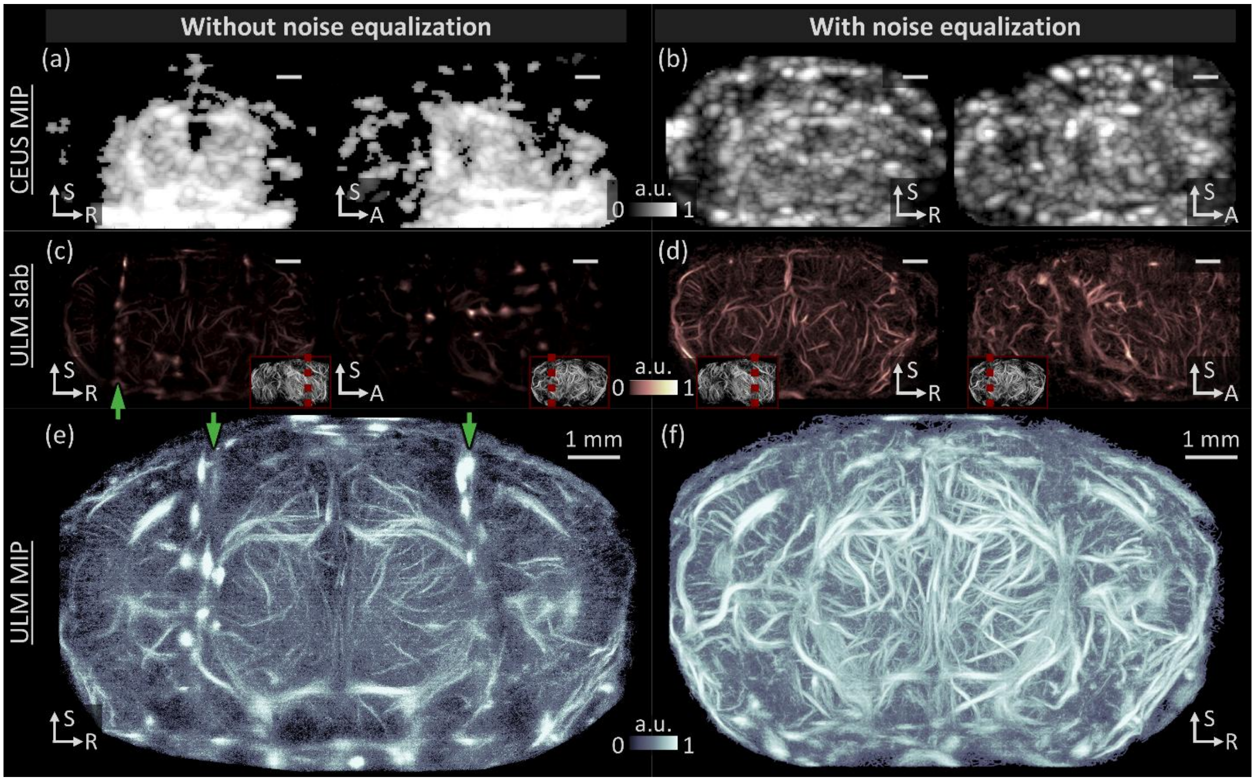


**Figure S1.** CEUS and ULM image quality comparison with noise equalization (right panels) and without noise equalization (left panels). Noise equalization improved image quality in both modalities. (a-b) MIP of 400 CEUS frames. (c-d) 2 mm slabs of ULM image constructed from 40,000 CEUS frames. The thumbnail in the red box indicates the slab location. The selected view cuts through regions with strong artifacts, and green arrows indicate artifact locations. In (a-d), all scale bars indicate 1 mm. A: anterior; R: right; S: superior.

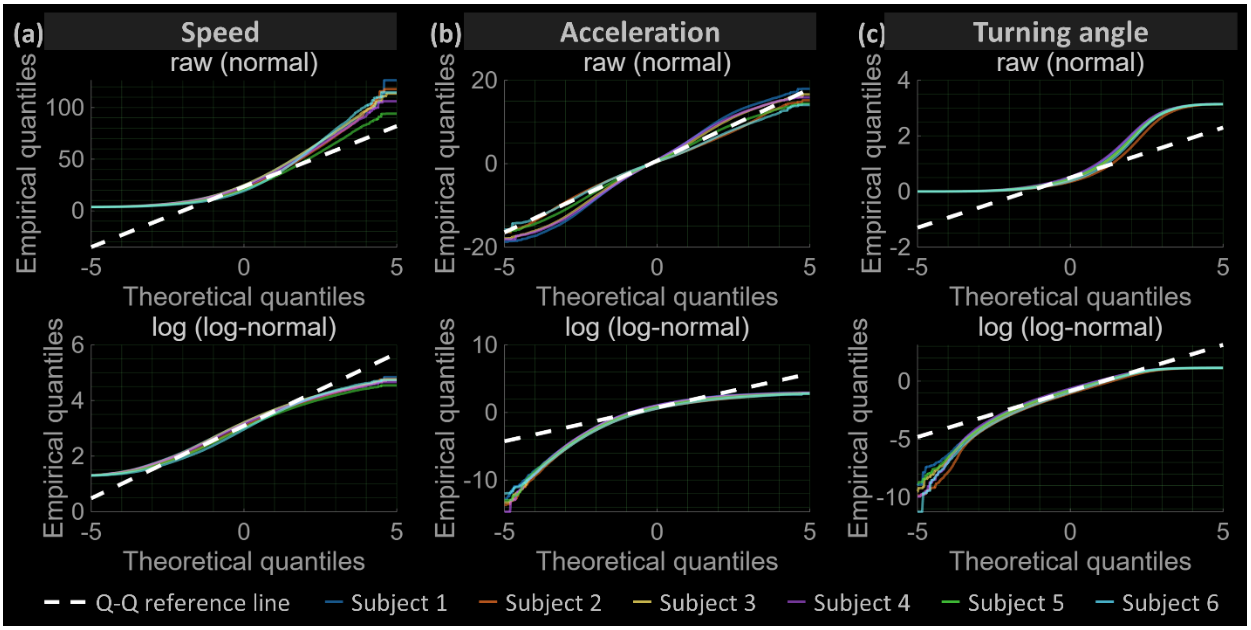


**Figure S2.** Q–Q plots assessing the distributions of speed (a), acceleration (b), and turning angle (c). Speed and turning angle are consistent with log-normal distributions, whereas acceleration is consistent with a normal distribution. The upper row evaluates normality using raw values, and the lower row evaluates log-normality using natural log-transformed values compared with theoretical quantiles. For acceleration, log-normality was assessed using the absolute values of the raw data. Colored lines represent individual subjects. The dashed white line indicates the IQR-fitted Q–Q reference line.

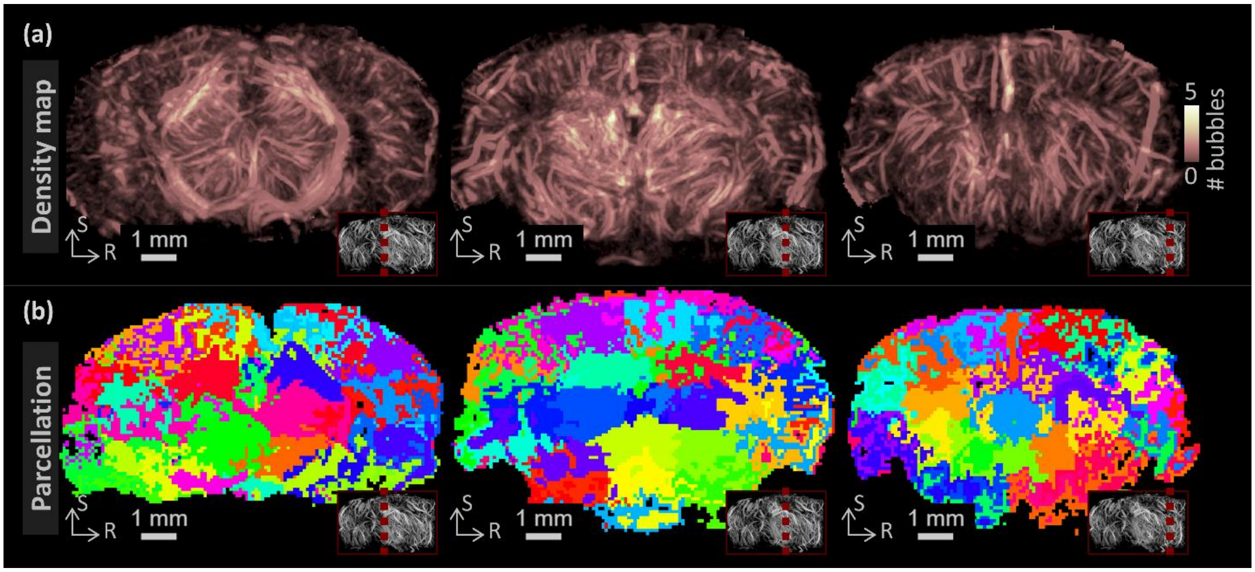


**Figure S3.** Group-wise density map (a) and its derived parcellation (b). The thumbnail in the red box indicates the slice location. In panel b, each color indicates a distinct parcel, unlabeled. R: right; S: superior.

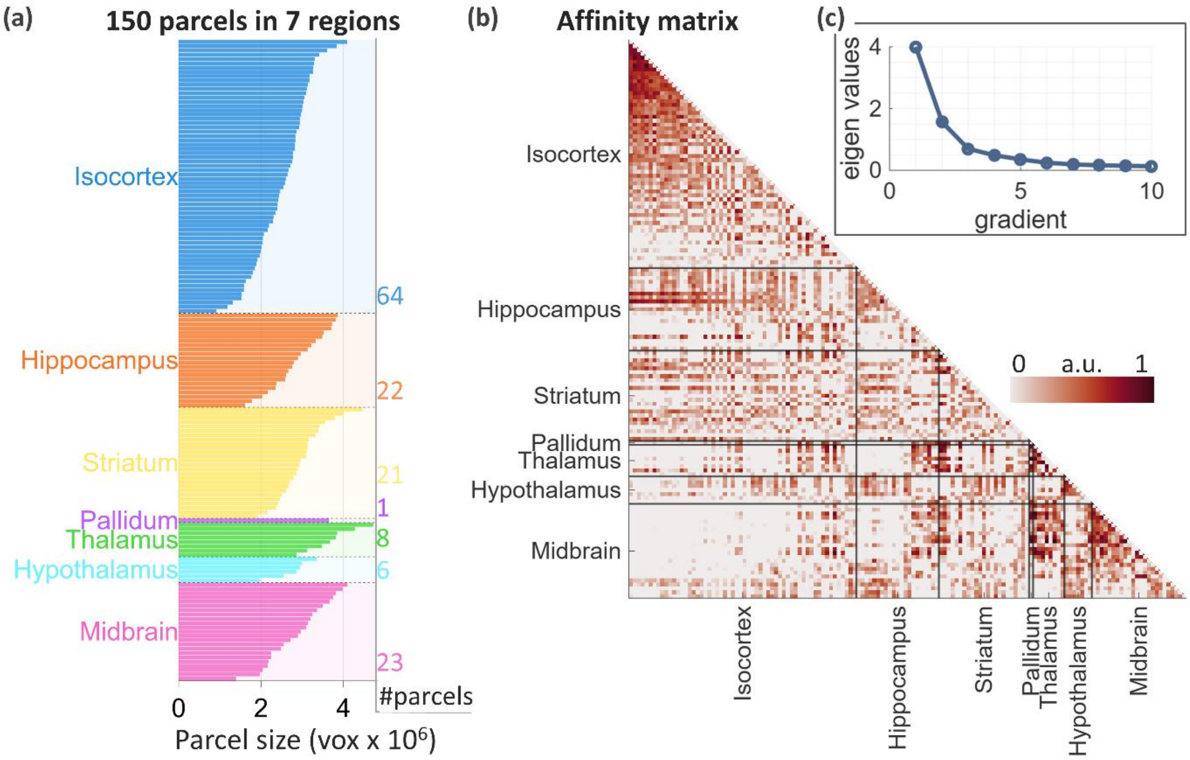


**Figure S4.** Group-consensus parcellation results summarized in seven brain regions (a), affinity matrix derived from these 150 brain parcels for diffusion embedding (b), and ranked eigenvalues of the diffusion operator, M, showing two dominant gradients (c).

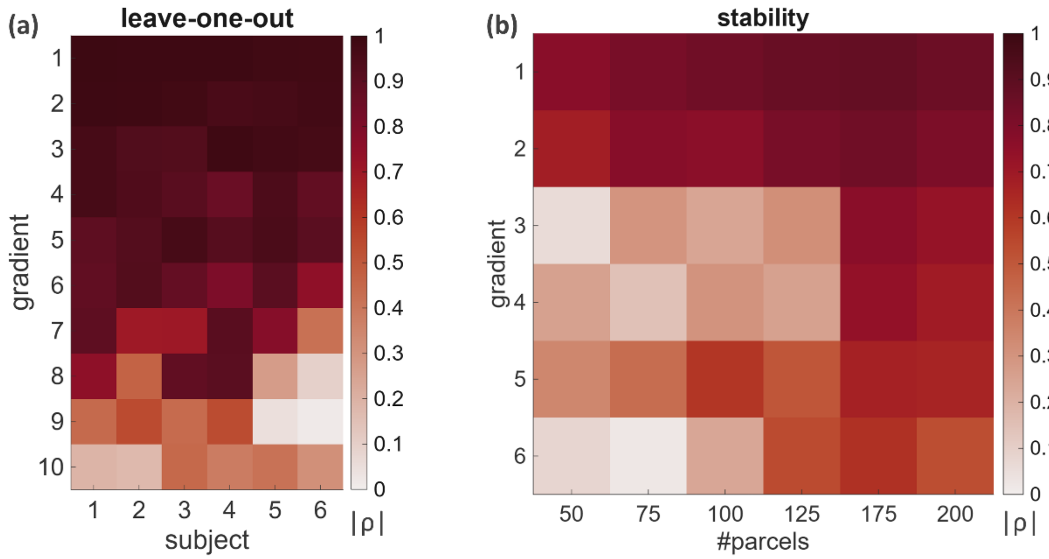


**Figure S5.** Gradient embedding validation showing the consistency of the top two gradients. (a) Pearson's ρ between the top ten gradients derived from the group affinity matrix and those derived from the affinity matrix from leave-one-subject-out. (b)

Pearson's ρ between the top six gradients derived based on 150 group-level parcels and those derived based on 50 to 200 group-level parcels.
